# RNA as a component of fibrils from Alzheimer’s disease and other neurodegenerations

**DOI:** 10.1101/2023.02.01.526613

**Authors:** Leslie R. Bridges

## Abstract

Fibrils from brains of patients with Alzheimer’s disease^1–5^, Parkinson’s disease^6^, amyotrophic lateral sclerosis^7^ and other neurodegenerations^3,4,8–18^ contain unknown molecules. Extra densities (EDs), containing these unknown molecules, are available to examine in electron cryo-microscopy maps from the Electron Microscopy Data Bank^19^, a public repository. EDs can be visualised in their protein environments using matched atomic models from the Protein Data Bank^20^, another public repository. Lysine-coordinating EDs from a wide range of neurodegenerative diseases^1–6,8–18^ and EDs from the glycine-rich region of TAR DNA-binding protein 43 (TDP-43) fibrils in amyotrophic lateral sclerosis with frontotemporal lobar degeneration (ALS-FTLD)^7^ were the subject of the present study. EDs ran parallel to the fibril axis and at right angles to protein with a repeat distance matching that of protein. They formed connections with protein consistent with a role in the guided assembly of fibrils. They had a connectivity pattern and estimated molecular weights consistent with ribonucleic acid (RNA). A straight form of RNA (ortho-RNA, oRNA) was modelled into one ED. It fitted other EDs and formed a rich symmetrical network of hydrogen bonds when docked to protein, implicating RNA as a unifying and organising factor in neurodegeneration. A new hypothesis of neurodegeneration (ponc, <u>p</u>rotein <u>o</u>rtho-<u>n</u>ucleic acid <u>c</u>omplex, pronounced ponk) is proposed in which RNA is the driver of these diseases. According to the ponc hypothesis, a particular RNA sequence (likely repetitive) enciphers a particular strain of ponc agent with its own protein fold and type of neurodegeneration. Ponc provides an explanation of fibril growth and replication, species barrier and adaptation, inherited neurodegeneration, resistance to chemicals and irradiation, protein-free transmission and co-pathologies. Ponc may also be relevant to other chronic diseases and origins of life. New treatments might be possible, targeting the unique chemical and physical properties of ponc.

---

Electron cryo-microscopy (cryo-EM) maps of fibrils from brains of patients with Alzheimer’s disease (AD)^1–5^ and other neurodegenerations^3,4,6–18^ contain EDs, whose constituent molecules are unknown. In one class of EDs, constituent molecules coordinate with one to four lysines of certain proteins, namely (i) alpha-synuclein (α-syn), in dementia with Lewy bodies (DLB)^6^, multiple system atrophy (MSA)^11^ and Parkinson’s disease (PD)^6^, (ii) tau, in AD^1–5^, argyrophilic grain disease (AGD)^13^, chronic traumatic encephalopathy (CTE)^9^, corticobasal degeneration (CBD)^3,10^, Gerstmann-Sträussler-Scheinker disease (GSS)^12^, globular glial tauopathy (GGT)^13^, Pick’s disease (PiD)^8^, primary age-related tauopathy (PART)^4^, progressive supranuclear palsy (PSP)^13,16^ and prion protein-cerebral amyloid angiopathy (PrP-CAA)^12^ (iii) PrP, in GSS^18^ and (iv) transmembrane protein 106B (TMEM106B)^14–17^, in diverse neurodegenerations and aging. Lysine-coordinating EDs^1–6,8–18^ and EDs from the glycine-rich region of TDP-43 fibrils in ALS-FTLD^7^ (hereinafter referred to simply as EDs) were the subject of the present study. The aim was to identify the constituent molecule of EDs, by *in silico* methods, using published cryo-EM and atomic data from the Electron Microscopy Data Bank (EMDB)^19^ and Protein Data Bank (PDB)^20^ public repositories.

## The nature of EDs

EDs run parallel to the fibril axis, consistent with a structural role as guide and support (Fig. 1). They run at right angles to and in register with protein strands (the repeat distance, about 4.8 Å, is the same, Extended Data Fig. 1a). They twist and arc with the fibril, maintaining a constant distance and attitude to the protein. Typically, they are continuous with distinct repeating features consistent with a straight polymer (Extended Data Fig. 1b). For example, ED 10650-43 in alpha-synucleinopathy MSA^11^ is an elliptic rod with height 10.1 Å, width 6.7 Å and aspect ratio 2:3 (EDs are named by the EMDB number and first coordinating residue). It has distinct features at the vertices and co-vertices (V1, V2, CV1 and CV2, Extended Data Fig. 1c-f) indicating direction, rotation and chirality. EDs are correlated and, allowing for differences in resolution and occupancy, their features match, consistent with a common constituent molecule (Extended Data Fig. 2). EDs have a three-blob connectivity pattern (Extended Data Fig. 2) consistent with RNA^21^ and inconsistent with protein, glycogen, heparin and poly (ADP-ribose). At or near full occupancy, estimated molecular weights are about 300 Da consistent with RNA (Extended Data Table 1). In EDs with weaker occupancy estimated molecular weights are correspondingly lower. In ED 3742-317 in AD^1^, the estimated molecular weight is about double that of RNA, raising the possibility of a duplex. The mass of an ED is about 1-2% of the mass of the fibril.

**Fig. 1.**
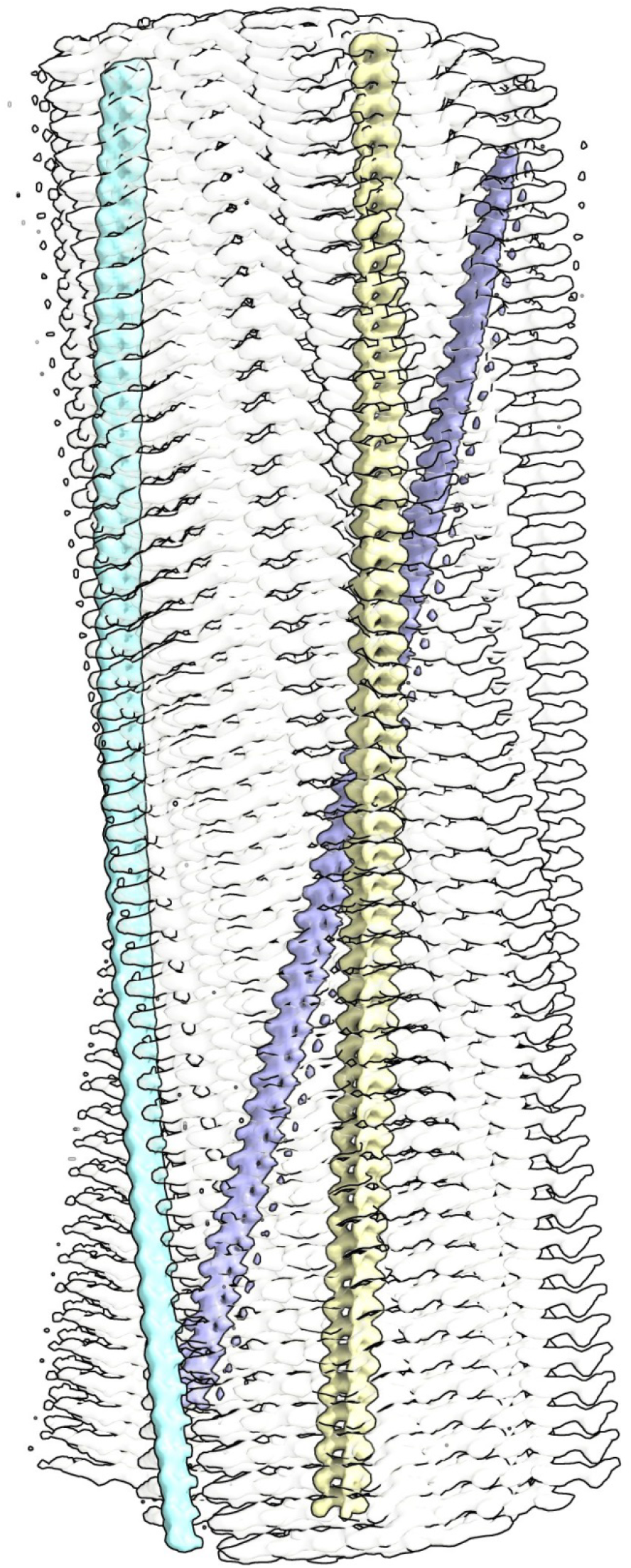
Guide and support. Alpha-synuclein fibril of multiple system atrophy. Extra densities (colours) are upright, at right angles to protein rungs (transparent, white), consistent with a structural role as guide and support. Distance between protein rungs is 4.7 Å. Image of EMDB 10650^11^ created with UCSF ChimeraX^45^.

## The protein environment of EDs

EDs are typically within hydrogen-bonding distance of protein. At authors’ recommended contour levels, EDs are mainly detached from protein but at reduced level (corresponding to increased map volume) specific connections (or tethering patterns) are seen, consistent with hydrogen bonds (Extended Data Fig. 3a). This suggests a role for EDs in the guided assembly of fibrils, by both guided folding and guided stacking, by the linking together of protein residues, chains and protofilaments. In alpha-synucleinopathies DLB and PD^6^, an ED coordinates with motifs 32**K**T**K**34 and 43**K**T**K**45 in a groove in the fibril (residues facing the ED are in bold). In alpha-synucleinopathy MSA^11^, an ED coordinates with 43**K**T**K**45 on opposing protofilaments in a cavity in the fibril (Extended Data Fig. 3a). Other EDs are found on the outside of the fibril at 32**K**T**K**34 and 58**K**T**K**60. In the tauopathies, EDs coordinate with the motif 317**K**V**T**S**K**321 in paired helical filaments (PHFs)^1,3,5,12^, straight filaments (SFs)^1,2,4,12^ and fibrils from CTE^9^, PiD^8^, PSP^13,16^ and GGT^13^. In PHFs^1,3,5,12^, an ED, arranged horizontally with respect to 317**K**V**T**S**K**321, forms a two-point tethering pattern with K317 and K321 on the outside of the fibril. In PSP with Richardson’s syndrome (PSP-RS)^13^, an ED, arranged horizontally, forms a three-point tethering pattern with K317, K321 and K340 in a cavity in the fibril (Extended Data Fig. 3b). In GGT^13^, an ED, arranged vertically, forms a four-point tethering pattern with K317, K321, N327 and K340 in a cavity in the fibril (Extended Data Fig. 3c). In straight filaments from AD^1,2,4^, PART^4^ and PrP-CAA^12^, an ED coordinates with K317 and K321 in a cleft between opposing protofilaments. The tethering patterns are similar, suggesting that the constituent molecules of EDs in different diseases may exert an equivalent tethering effect. EDs coordinate with 290**K**CG**SK**294 in CBD^3,10^ and 290**K**C**G**S**K**294 in AGD^13^, both in cavities in the fibrils. In CBD^3,10^, the ED coordinates with K290, K294 and K370 (Extended Data Fig. 3d). This is a hydrogen-bonding donor atom environment seeking acceptor atoms in the constituent molecule. In AGD^13^, the ED coordinates additionally with E372 (Extended Data Fig. 3e). Since glutamate has hydrogen-bonding acceptor atoms, it seeks donor atoms in the constituent molecule. It is reasonable to infer that constituent molecules and their protein environments are matched by their steric and hydrogen-bonding properties. In GSS^18^, EDs coordinate with 104**K**P**K**106 and 110**KH**111 of PrP on the outside of the fibril. In fibrils of TMEM106B, an ED coordinates with 178**K**A**R**180^14–17^ in the gap between opposing protofilaments. In ALS-FTLD^7^, EDs coordinate with glycine-rich (non-lysine) environments at 303**QG**304 and 306**N**M**G**308 of TDP-43 on the outside of the fibril.

## The constituent molecule of EDs

Connectivity and molecular weight point to RNA (see above). The continuous repeating nature of EDs suggests a polymer, whilst the protein environment suggests a polyanion, with abundant hydrogen-bonding acceptor atoms and fewer donor atoms consistent with RNA. EDs are axially continuous with distinct repeating features, consistent with a polymer in step with the helical rise. Ions^22^ and protein tails (whether 7EFE9 from tau^1^ or 75GG76 from ubiquitin^3^) would be axially discontinuous (or non-distinctive and non-repeating, if out of step with the helical rise). A straight form of RNA with 4.8 Å repeat has been described in RNA-tau fibrils grown *in vitro*^21^. Furthermore, a straight form of nucleic acid was postulated by Watson and Crick^23^. Fibril tails in scrapie-associated fibrils (SAF) were queried as RNA^24^.

Mechanically disrupted tau fibrils appear to be joined by a thread^25^. RNA is present in plaques, tangles and other neurodegenerative lesions^26^. Due to map averaging, it was not possible to read the RNA sequence directly in the present study, but since the protein interface is identical at every rung, it seems likely, for consistent binding, that the RNA is repetitive. This could be as a homopolymer or short sequence repeat. Earlier studies are against a foreign nucleic acid^27^, but host-like repetitive sequences are present^28^, variable lengths are possible^29^, and a single-stranded deoxyribonucleic acid (ssDNA) with a repeat palindromic sequence (TACGTA)n was found in scrapie^30^. Sense and antisense transcripts of short sequence repeats are found in many genetic neurodegenerations, including Huntington’s disease (CAG/CTG), C9orf72 ALS-FTLD (GGGGCC/CCCCGG) and Friedreich’s ataxia (GAA/TTC)^31^. Experimentally, nucleic acid promotes fibrillization of PrP^32^, tau^33^ and alpha-synuclein^34^. Some RNA sequences are more potent at this than others^32^, consistent with preferential (non-exclusive) binding to the target protein.

## Ortho-RNA (oRNA)

RNA (polyU) was modelled into ED 10650-43^11^. In Molprobity^35^, it has an unrecognised straight backbone but otherwise perfect geometry (clash score 0, bad sugar puckers 0, bad bonds 0 and bad angles 0). The name ortho-RNA is proposed in view of its straight (ortho) form and orthogonal pose relative to protein. oRNA is stabilised by base stacking, a hydrogen bond between HO2’ and O2 and carbon hydrogen bonds between H2’ and O3’ and H6 and O4’ (Extended Data Fig. 4a). There are unfavourable negative-negative interactions between O1P and O2P and acceptor-acceptor interactions between O1P and O5’ but these are conceivably offset in the protein environment. oRNA has a three-blob connectivity pattern^21^(Extended Data Fig. 4b), the same as EDs (Extended Data Fig. 2). Two phosphorus atoms and the distal carbon atom C1 form a near-orthogonal triangle (Extended Data Fig. 4c), offering a feasible basis for orthogonal lattice formation with protein. Phosphorus-phosphorus distances of 4.713, 4.781, 4.801 and 4.842 Å were modelled and combinable, allowing for some flexibility. oRNA fits ED 10650-43^11^ and other EDs, consistent with a unifying role in these diseases (Fig. 2). Hydrogen-bonding atoms mapped to the surface of oRNAs show base-dependent patterns resembling bar codes (Extended Data Fig. 5). It is proposed that these patterns are recognised by regions of the target protein, where particular oRNA sequences are preferred. Other versions of the model, also with good geometry, were as (i) any base sequence, (ii) ssDNA, (iii) abasic RNA and (iv) a duplex. The duplex (not the same as double-stranded RNA, dsRNA) was made by docking together two molecules of oRNA (Extended Data Fig. 4d). Any base sequence is permitted in both strands and there is a rich network of hydrogen bonds between whole nucleotides (not Watson-Crick in type). Unlike dsRNA, it is not mandated for the strands to be antiparallel and complementary.

**Fig. 2.**
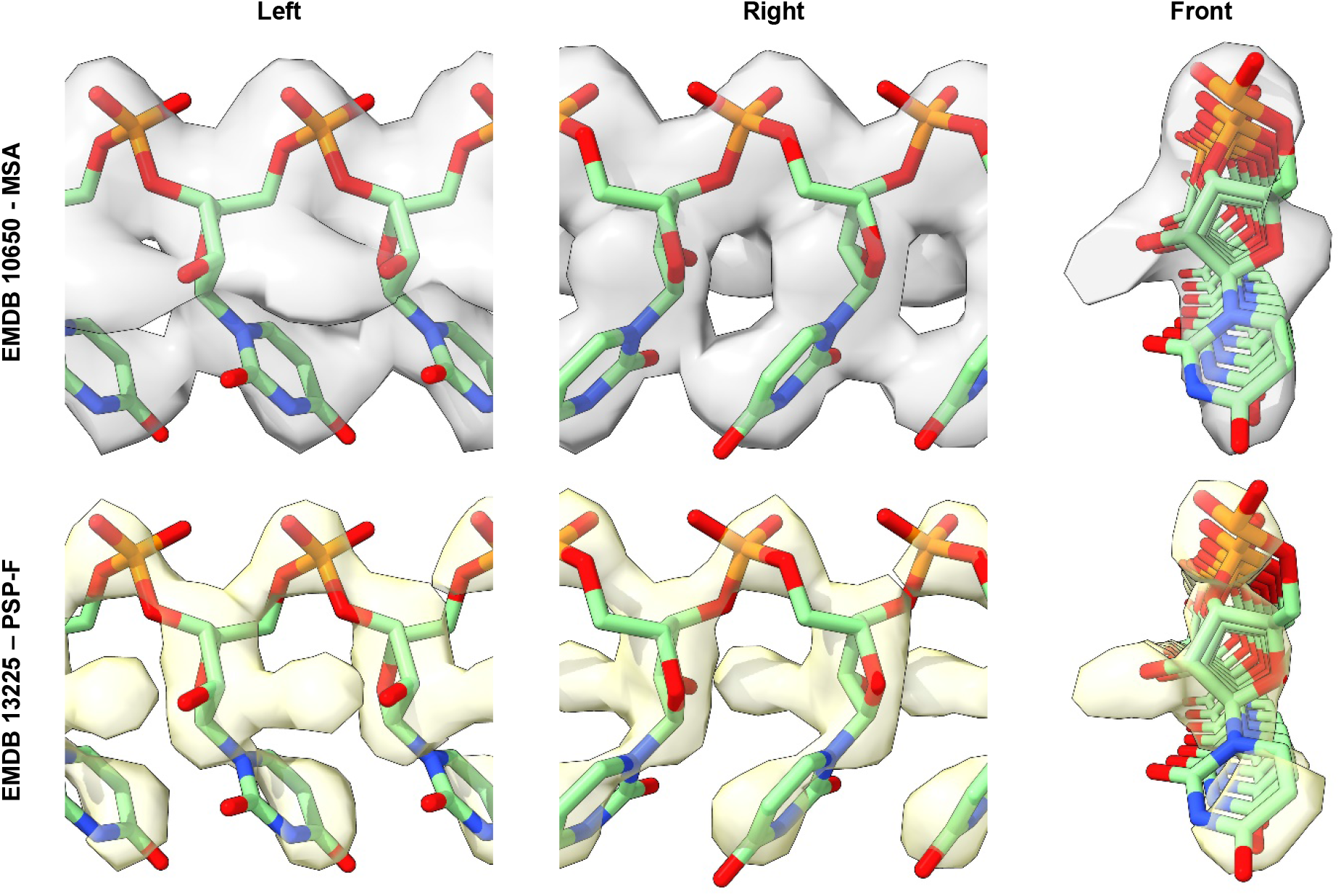
Unifying factor. Ortho-RNA fits extra densities from alpha-synuclein fibril of multiple system atrophy and tau fibril of progressive supranuclear palsy with frontal presentation with comparable fidelity, consistent with a unifying role in these diseases. Images of EMDB 10650^11^ and EMDB 13225^13^ created with UCSF ChimeraX^45^.

## Molecular docking of oRNA to protein

Docked poses of oRNA to protein show a rich symmetrical network of hydrogen bonds, favourable AutoDock Vina scores^36^, favourable Molprobity clashscores^35^ and overlap with EDs, consistent with an organising role in neurodegeneration (Fig. 3, Extended Data Fig. 6, Extended Data Table 1). This is true at the glycine-rich region of TDP-43 fibrils in ALS-FTLD (Extended Data Fig 6d), as well as lysine-containing environments in other cases. Docking of oRNAs polyU, polyC, polyA and polyG in AGD indicates base-dependency of tethering patterns (Extended Data Fig. 7), consistent with RNA sequence as a determinant of fibril structure. By analogy to 110**KH**111 in PrP fibrils^18^ (Extended Data Fig. 6c), oRNA is dockable to a similar motif, 14**H**Q**K**16, in beta-amyloid (Abeta) fibrils^37^ (Extended Data Fig. 6h), even though the map, EMDB 13800, lacks an ED at this site. Similarly, one of the published maps of CTE^9^ (EMDB 0528) has an ED at 317**K**V**T**S**K**321, whereas the other map (EMDB 0527) lacks an ED at this site. Conceivably, fibrils, by retaining an imprint, can lose and regain oRNA. oRNA is also dockable to 317**K**V**T**S**K**321 of the tau fibril of PiD^8^ (Extended Data Fig. 6i; the map, EMDB 0077, shows no ED, but the paper mentions an ED at this site). There are currently no cryo-EM maps of amyloid-Bri (Abri) in familial British dementia or amyloid-Dan (ADan) in familial Danish dementia, but inspection of their protein sequences^38^ suggests possible binding sites for oRNA at 27**KK**28 in Abri and 32**KH**33 in ADan. Electrostatic potential maps suggest that electronegative oRNA and electropositive protein bind together as opposites (Extended Data Fig. 8). The separation of charges indicates an electric dipole, raising the possibility of a novel therapeutic strategy for the disruption of fibrils, by the application of external electric or magnetic fields. Duplex oRNA is dockable within double-sized ED 3742-317 in PHF tau fibril in AD^1^ (Extended Data Fig. 6g). Docking poses for ortho-ssDNA were found, similar to those of oRNA, suggesting that ssDNA may substitute for RNA in some fibrils. In a few cases, AutoDock Vina and Molprobity scores for oRNA were unfavourable. This might suggest a methodological issue addressable with flexible protein docking in future studies, or a smaller constituent molecule such as ssDNA or abasic RNA in some cases.

**Fig. 3.**
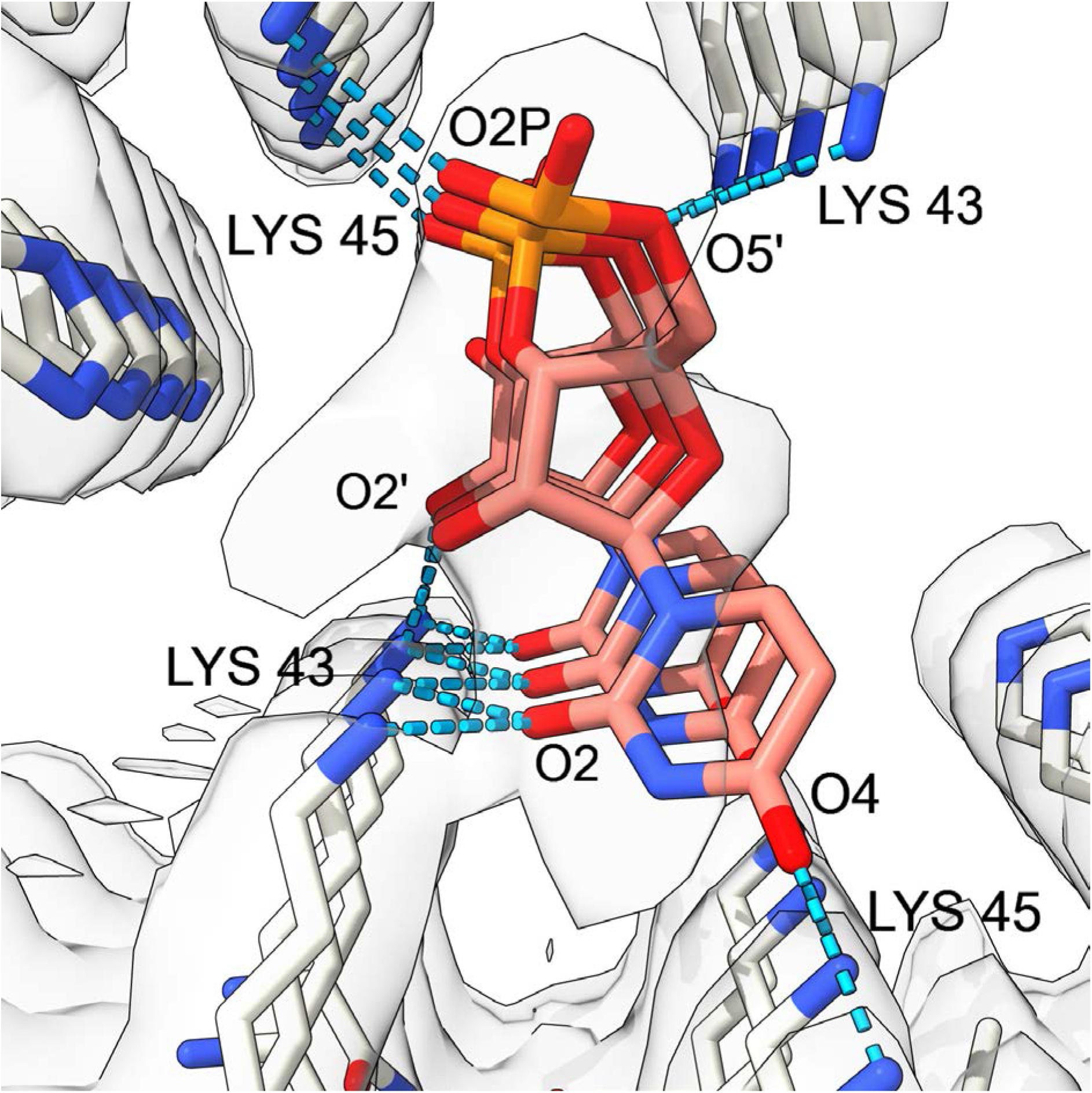
Organising factor. Ortho-RNA in extra density of alpha-synuclein fibril of multiple system atrophy, linked to protein by a rich symmetrical network of hydrogen bonds, consistent with an organising role in this disease. Image of EMDB 10650 and PDB 6xyo^11^ created with UCSF ChimeraX^45^, oRNA pose generated with AutoDock Vina^36^.

## Ponc, a new hypothesis of neurodegeneration

A new hypothesis of neurodegeneration is proposed, namely ponc (protein ortho-nucleic acid complex, pronounced ponk). Like the virino^39^, holoprion^40^ and nemavirus^30^ hypotheses, but unlike the prion hypothesis^41^, ponc affirms the presence of nucleic acid in the particles that cause neurodegeneration. Based on the evidence herein, the following is proposed:

i. Ponc arises from the interaction of a particular sequence of RNA with a particular sequence of protein to form a complex (rogue RNA meets rogue protein).
ii. The RNA sequence enciphers the essential transmissible component (or strain), but the disease properties (including the protein conformation) are co-dependent on the protein sequence in the host. Since the protein is from the host, there is no requirement for the RNA to be translated.
iii. Ponc is resistant to chemicals and irradiation because oRNA is braced by protein and the RNA sequence is repetitive and of variable length.
iv. A species barrier arises when ponc RNA encounters a new target protein, in a new host, with reduced affinity. Adaptation is due to replacement of the RNA with a sequence better-matched to the new target protein.
v. Inherited disease occurs when a particular RNA encounters a new target protein, with increased affinity, due to mutation of the protein in the host.
vi. Protein-free transmission is due to transfer of RNA.
vii. Co-pathologies arise where an RNA and its complement have more than one target protein. For example, in AD it is envisaged that complementary RNA from an Abeta ponc interacts with tau to form a tau ponc and *vice versa*. This suggests a bidirectional (symbiotic) relationship, in contradistinction to the unidirectional relationship of the amyloid cascade hypothesis^42^.
viii. Ponc-inducing RNA could be endogenous (in certain cells in certain states and cumulative with age) or exogenous (from injection or ingestion).
ix. Self-annealing, repetitive nucleic acid might act as a lasso, entangling mitochondrial DNA to form multimers^30^ (mitochondrial injury is a known pathogenetic factor in neurodegeneration).
x. Ponc might be the cause of other chronic diseases, such as amyloidosis and inclusion body myositis (RNA and PHF tau fibrils are present in diseased muscle in IBM^43^).

Ponc is a testable hypothesis. Repetitive RNAs and neurodegenerative proteins could be mixed *in vitro* and *in silico* to determine structural changes and free energies of binding. Repetitive RNAs could also be introduced into cell lines and experimental animals. Conceivably, some repetitive RNAs might be a biohazard.

## Ponc and origins of life

Observations here on tethering patterns and docking, suggest that a nucleotide of oRNA can link to consecutive protein rungs and *vice versa* (Extended Data Fig. 9a,b). This suggests a mechanism for fibril growth whereby, iteratively, the leading nucleotide forms a hydrogen bond with a new protein rung and the leading protein rung forms a hydrogen bond with a new nucleotide (Extended Data Fig. 9c). Ponc itself might be a catalyst/ribozyme/enzyme for hybridization and ligation of nucleotides (oRNA polymerase) and guided stacking of protein (stackase). Similarly, ponc might function as a protein foldase, via specific tethering of protein by oRNA (Extended Data Fig. 9d). Evidence here for a duplex form of oRNA raises the possibility of RNA replication (Extended Data Fig. 9e). Steric factors prohibit Watson-Crick binding to oRNA, but regions of oRNA might unfold to normal RNA, replicate to dsRNA and refold as duplex oRNA. Ponc itself might be a catalyst/ribozyme/enzyme for unfolding/refolding and nucleotide hybridization and ligation of the complementary strand (oRNA replicase). Ponc, similar to a virus, might use host cellular resources (nucleotides, proteins and ATP) for its growth and replication. Experimentally, *in vitro*, nucleic acids and short basic peptides such as (KL)3 form complexes, and an amyloid-nucleic acid (ANA) world has been proposed^44^ analogous to the RNA world. Ponc offers a specific structural basis for an ANA world. The ponc protein interface, like a *tabula rasa*, might generate new RNAs, with structural or catalytic properties favourable for survival, and form new poncs, all working together in the primordial soup. Storage of genetic information by a protein interface, in ponc, is a departure from known genetic systems. RNA and protein in ponc, whilst co-dependent, are not equal: although RNA is the lesser component by mass, it has primacy because it replicates, whereas protein is merely enriched. Ponc can be compared to the DNA double helix. Both are helical and both generate and store biological information, but pairing is exclusive in DNA and likely non-exclusive in ponc. Ponc can be seen as an emergent property of matter, an autonomous entity capable of growth and replication, the product of a molecular serendipity (the distance 4.8 Å, the common spacing of its nucleic acid and protein components), and a form of life, but not as we know it. Given the unique chemical and physical properties of ponc, different to those of other cellular structures, it might be possible to destroy it, leaving the cell intact, raising the possibility of future new treatments for Alzheimer’s disease, Parkinson’s disease, ALS and other neurodegenerations.

## Methods

### Data sources and selection

Data for this study were sourced from public repositories, the EMDB^19^ and the PDB^20^. Cryo-EM maps and protein models of fibrils from autopsy brains of patients with the following neurodegenerative diseases were examined: AD^1–5,15,37,46^, AGD^13^, ALS-FTLD^7^, CTE^9^, CBD^3,10^, DLB^6^, FTLD-TDP^14,16^, GSS^12,18^, GGT^13^, MSA^11,15^, PD^6,17^, PiD^8^, PART^4^, PSP^13,16^ and PrP-CAA^12^. The proteins involved were Abeta^37,46^, alpha-synuclein^6,11^, PrP^18^, tau^1–5,8–10,12,13,16^, TDP-43^7^ and TMEM106B^14–17^. A majority of fibrils had EDs, as determined by inspection and reference to the published papers. Lysine-coordinating EDs^1–6,8–18^ (the commonest type of EDs) and EDs from the glycine-rich region of TDP-43 fibrils^7^ (which had a similar morphology) were the subject of the present study. EDs from other environments (*e.g*. the hydrophobic cavity in tau fibrils from CTE^9^) were excluded. EDs were named by the EMDB number and the first coordinating residue. Where pairs of EDs were present in doublets, associated protofilaments were denoted “a” and “b” (left and right respectively, in the opening position). Observations and measurements were made on the following EDs: 10650-32, 10650-43, 10650-58, 10651-32, 10651-43 and 10652-43 in alpha-synuclein fibrils in MSA^11^; 10512-290, 10512-343, 10514-290a and 10514-290b from tau fibrils in CBD^10^; 21200-290a, 21200-290b and 21201-290 in tau fibrils in CBD^3^; 21207-317a and 21207-317b in tau fibril in AD^3^; 0528-317a and 0528-317b in tau fibrils in CTE^9^; 0260-317 in tau fibril in AD^2^; 3741-317a, 3741-317b, 3742-317a, 3742-317b and 3743-317 in tau fibrils in AD^1^; 12550-317 in tau fibril in PART^4^; 12552-317 in tau fibril in AD^4^; 13218-317 and 13218-280 in tau fibril in PSP-RS^13^; 13219-317, 13219-280, 13220-317a, 13220-317b, 13220-290a, 13220-290b, 13221-317a and 13221-317b in tau fibrils in GGT^13^; 13223-317, 13223-280, 13224-317, 13224-280, 13225-317a, 13225-317b, 13225-280a and 13225-280b in tau fibrils in PSP-F^13^; 13226-290, 13226-281, 13227-290a and 13227-290b in tau fibrils in AGD^13^; 23890-317, 23871-317a and 23871-317b in tau fibrils in PrP-CAA^12^; 23894-317a and 23894-317b in tau fibrils in GSS^12^; 13708-303, 13708-306, 13710-303, 13710-306, 13712-303 and 13712-306 in TDP-43 fibrils in ALS-FTLD^7^; 14176-178 in TMEM106B fibril in MSA^15^; 24954-180 in TMEM106B fibril in FTLD-TDP^14^; 15285-32 from alpha-synuclein fibril in DLB and PD^6^; 26613-104a, 26613-104b, 26613-110a, 26613-110b, 26607-104a, 26607-104b, 26607-110a and 26607-110b in PrP fibrils in GSS^18^; 26663-317a, 26663-317b, 26664-317a, 26664-317b, 26665-317a and 26665-317b from tau fibrils in AD^5^.

### Observations and measurements

Observations and measurements were made in UCSF ChimeraX^45^ unless otherwise stated. A Dell Desktop-GUQ6HDT XPS 15 9500 with Intel^®^ Core™ i7-10750H CPU @2.6 GHz with 16 GB RAM running Windows 11 Home (version 21H2) with broadband internet connection was used to run software and to access data and software online. Each cryo-EM map was examined at the authors’ recommended contour level with the protein model *in situ*.

For clarity, the map was made transparent and clipped to the region of the protein. EDs were identified in their protein environment, by inspection and reference to the published papers. Allowing for differences in resolution and occupancy, EDs from the subject group were similar (*e.g*. they were straight, continuous, repeating and parallel to the fibril axis). To determine whether they were in perfect register with the protein, markers were placed 30 units apart on EDs and 30 rungs apart on the adjacent protein density and the distances compared. To determine whether EDs maintained a constant distance and attitude to the protein, slices of the fibril, 5 Å in thickness and 100 Å apart, were compared with the fitmap command. To assess whether EDs were within hydrogen-bonding distance of protein, color zones from the NZ (terminal nitrogen) atoms of lysines were incremented until they reached the surface of the EDs.

In well-resolved cases, EDs were detached from, or showed minor regions of fusion to, the protein density. At lower resolutions, some EDs were extensively fused to the protein density. EDs were dissected away from the rest of the density, in order to observe and measure them freely. This was done with the map eraser tool, by erasing the ED and subtracting the remainder from a copy of the original. Where there was extensive fusion between the ED and the protein density, the ED was isolated by using the color zone and split map commands, at a radius of 2 Å from the protein atoms. ED 10650-43^11^ was chosen as a reference ED, for comparison with other EDs, as it has relatively high resolution (2.6 Å) and strong occupancy. It is an elliptic rod with distinctive features at the vertices and co-vertices (Extended Data Fig. 1c-f). Viewed from the sides, in the authors’ opening orientation, it resembles a series of backward-facing chevrons. Further detail was evinced by adjusting the contour level. For example, at increased level V1 is joined by persistent density whereas V2 is no longer joined (Extended Data Fig. 2). Using such cues, EDs were oriented and compared to the reference ED. The height and width of EDs were measured using the measure blob tool, or by measuring between manually-placed markers. Correlation was measured by fitting 50 Å lengths of ED into the reference ED. After an initial manual placement, the fitmap command was invoked. The correlation metric in UCSF ChimeraX^45^ ranges from 0 to 1 (1 is a perfect fit). Colour-shading of repeating units was performed with Segger^47^ in UCSF ChimeraX^45^.

To examine connections between the ED and protein density, two copies of the cryo-EM map were viewed, superimposed. One copy was viewed opaque, at the authors’ contour level, and the other copy was viewed transparent, at reduced levels (as in Extended Data Fig. 3a). By adjusting the level incrementally, connections between the ED and protein density were determined in sequence. The connections were named tethering patterns. By reference to the atomic model, it was noted which protein residues formed part of the tethering pattern, whether the ED was vertical or horizontal with respect to the protein motif, and whether the ED was on the outside of the fibril or within a groove, cleft or cavity.

The molecular weight (MW) of the ED was calibrated to the MW of the protein. EDs were assessed as having high (or strong or near-stoichiometric) occupancy where, at the authors’ contour level, the volume was comparable to adjacent protein density. EDs were assessed as having low (or weak) occupancy where the volume was attenuated in comparison to the adjacent protein density. Estimates of MW were considered more representative for EDs with high occupancy. Protein density was isolated by removing non-protein density with the map eraser tool or, where protein and non-protein density were extensively fused, separation was carried out with the color zone and split map commands, with a 2 Å radius around protein atoms. Volumes of isolated EDs and protein densities were measured with the measure blobs tool or measure volume command. The MW of one rung of protein was calculated by opening the protein model in UCSF Chimera^48^, adding hydrogens, selecting one rung and invoking the keyboard shortcut ac mw. The ratio of the volume of ED to the volume of protein density was calculated. The ratio was multiplied by the MW of one rung of protein to give an estimate of the MW of one residue of ED. For comparison, the MW of RNA residues, in Daltons, are as follows: cytosine 304.2, uracil 305.2, adenine 328.2 and guanine 344.2.

### The model (ortho-RNA, oRNA)

A monomer with short phosphorus-phosphorus distance was selected from PDB 467d (B-DNA)^49^. In UCSF Chimera^48^, it was rebuilt as RNA (U) and torsions were adjusted for optimal fit in ED 10650-43^11^. A 60mer was built by moving monomers 4.7 Å apart (the distance between protein rungs in PDB 6xyo, the protein model corresponding to ED 10650-43). The 60mer was minimized in VMD^50^(**http://www.ks.uiuc.edu/Research/vmd/**) and NAMD^51^(**http://www.ks.uiuc.edu/Research/namd/**). A 7mer from the middle of the minimized 60mer was refined in Phenix.real_space_refine^52^, within the region of EMDB 10650 corresponding to ED 10650-43. The resulting 7mer was submitted to the MolProbity^35^ webserver. It showed a novel straight backbone and otherwise perfect geometry (clashscore 0, bad sugar puckers 0, bad bonds 0 and bad angles 0). The 7mer had residues with differing phosphorus-phosphorus distances. Individual residues were rebuilt in UCSF Chimera^48^ as four new 7mers with uniform spacings (4.713 Å, 4.781 Å, 4.801 Å and 4.842 Å). Further models were built with other single bases (A, C and G) and short sequence repeats (*e.g*. CAG and CUG). Other models were built as ssDNA and abasic RNA. All of the derived models showed the same as the original, namely a unique straight backbone and otherwise perfect geometry. Non-bonded interactions were examined in BIOVIA, Dassault Systèmes, Discovery Studio Visualizer, v21.1.0.20298, San Diego, Dassault Systèmes, 2021. This software was also used to depict hydrogen bonding surfaces for different bases. Features such as straightness, orthogonality and connectivity were examined in UCSF ChimeraX^45^. oRNA was fitted to EDs using a map simulated from the atoms at the same resolution as the ED (the correlation metric in UCSF ChimeraX^45^ was noted). Duplex models were created by designating a 7mer as receptor and a 3mer as ligand. They were converted to PDBQT format in AutoDockTools^53^. Molecular docking was done with AutoDock Vina^36^. The resulting models were examined in UCSF ChimeraX^45^. Hydrogen bonds were displayed with relaxed criteria (distance tolerance 0.4 Å, angle tolerance 20°). Duplexes were validated in the MolProbity Webserver^35^.

### Molecular docking

A 3mer of oRNA (UUU) was used as ligand. It was chosen with an inter-residue distance as close as possible to the inter-strand distance of the protein. Five rungs of protein (cut down to coordinating and intervening residues) were used as receptor. Where the PDB had less than five rungs, extra rungs were built it UCSF Chimera^48^ and validated in the MolProbity Webserver^35^. Ligand and receptor were converted to PDBQT format in AutoDockTools^53^. Both ligand and receptor were docked as rigid objects. Molecular docking was done with AutoDock Vina^36^. The resulting poses were examined in UCSF ChimeraX^45^. They were assessed for overlap with the ED, symmetrical hydrogen bonds (displayed with relaxed criteria as above) and favourable AutoDock Vina score (affinity, kcal/mol). The protein and selected pose were combined in UCSF Chimera^48^ and submitted to the MolProbity Webserver^35^ to determine the clashscore. Electrostatic potential surfaces were created in UCSF ChimeraX^45^, using the coulombic command, from models of oRNA docked to protein (the 3mer was extended to a 7mer for display purposes).

### Ethical statement

The cryo-EM maps and atomic models used in the present study were sourced from the EMDB and PDB public repositories, from the publications recorded herein, which all affirm that ethical review and informed consent were obtained and provide the names of the relevant organizations. All patient data for the present study was at one remove and no new patient data was obtained.

## Data availability

The EMDB and PDB accession numbers for data examined in this paper are provided in the text. Any other relevant data are available from the corresponding author upon request.

## Acknowledgements

The present work was done on cryo-EM and atomic models from the EMDB and PDB public repositories. The patients and relatives were thanked in the original publications and, at one remove, I also thank them. I also thank the scientists, software developers, data managers and funders whose efforts, at one remove, have made my own work possible. Molecular graphics and analyses performed with UCSF Chimera and UCSF ChimeraX were developed by the Resource for Biocomputing, Visualization, and Informatics at the University of California, San Francisco, UCSF Chimera with support from NIH P41-GM103311 and UCSF ChimeraX with support from National Institutes of Health R01-GM129325 and the Office of Cyber Infrastructure and Computational Biology, National Institute of Allergy and Infectious Diseases. VMD and NAMD were developed by the Theoretical and Computational Biophysics Group in the Beckman Institute for Advanced Science and Technology at the University of Illinois at Urbana-Champaign. I thank St George’s University Hospitals NHS Foundation Trust NHS and South West London Pathology for my employment and St George’s University of London for my honorary academic status.

## Author contribution

LRB performed the whole of this work.

---

**Extended Data Fig. 1.**
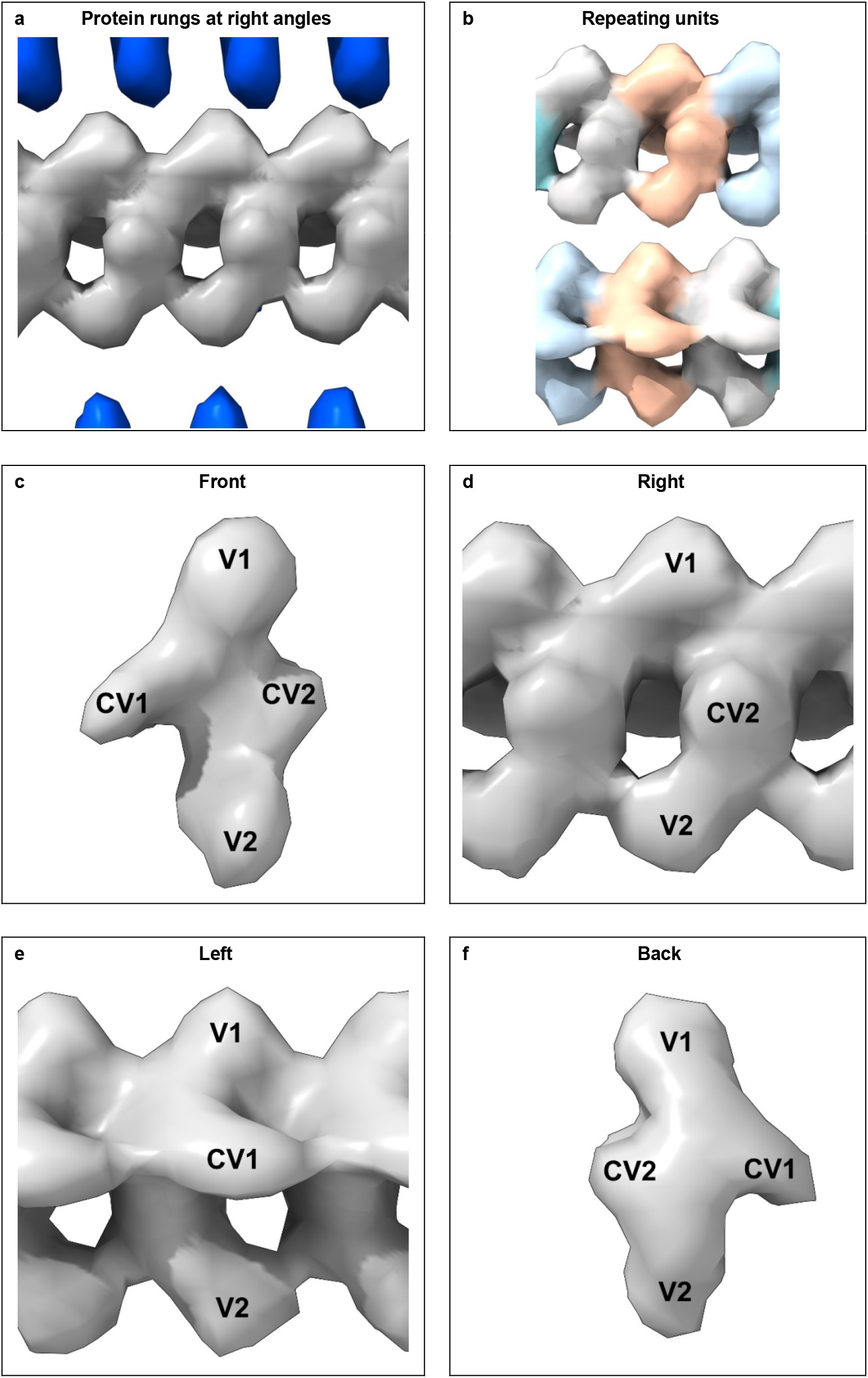
Straight polymer. Extra density from alpha-synuclein fibril of multiple system atrophy. Repeating features are consistent with a polymer. Images of EMDB 10650^11^ created with UCSF ChimeraX^45^, colour-shading of units with Segger^47^.

**Extended Data Fig. 2.**
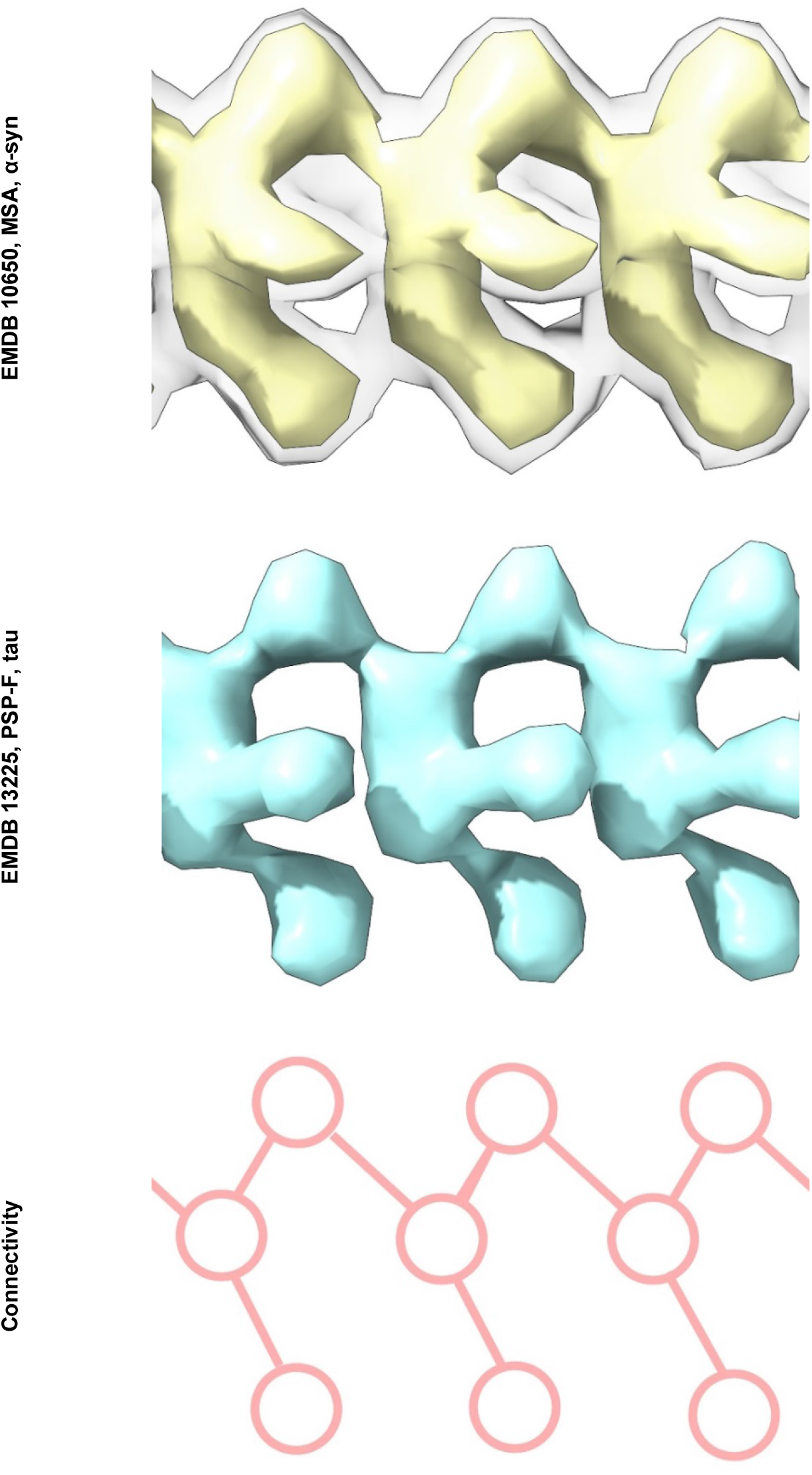
Common constituent molecule. Extra densities from alpha-synuclein fibril of multiple system atrophy and tau fibril of progressive supranuclear palsy with frontal presentation have features and connectivity consistent with a common constituent molecule. Images of EMDB 10650^11^ at two contour levels and EMDB 13225^13^ created with UCSF ChimeraX^45^, connectivity diagram created with ImageJ^54^.

**Extended Data Fig. 3.**
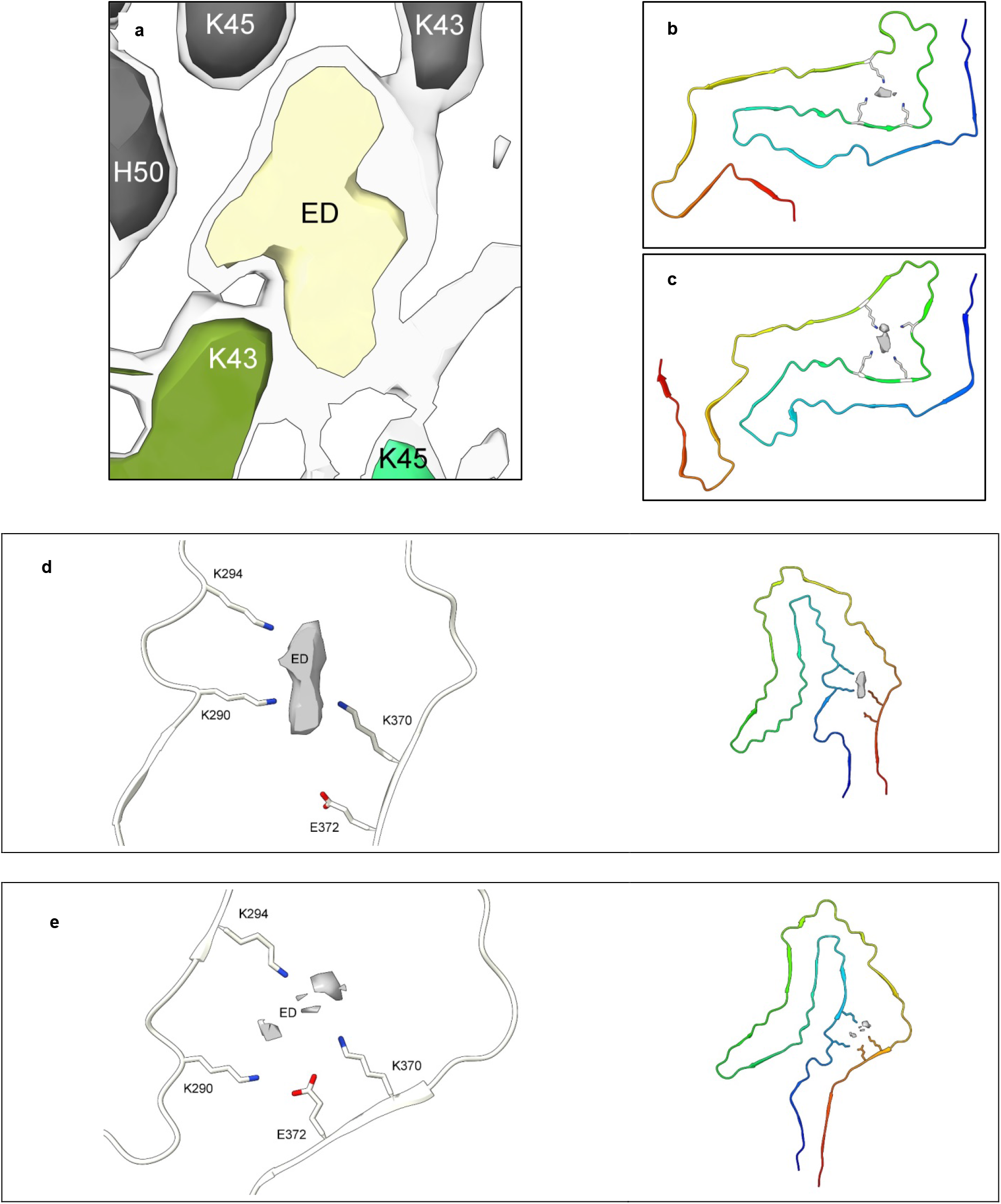
Tethering patterns. **a.** ED in α-syn fibril of MSA at two contour levels. **b-e.** EDs and protein models from tau fibrils of PSP-RS **(b)**, GGT **(c)**, CBD **(d)** and AGD **(e)**. Note specific connections or tethering patterns between EDs and protein. Images of EMDB 10650^11^ **(a)**, EMDB 13218 and PDB 7p65^13^ **(b)**, EMDB 13219 and PDB 7p66^13^ **(c)**, EMDB 10514 and PDB 6tjx^10^ **(d)**, EMDB 13226 and PDB 7p6d^13^ **(e)**, created with UCSF ChimeraX^45^.

**Extended Data Fig. 4.**
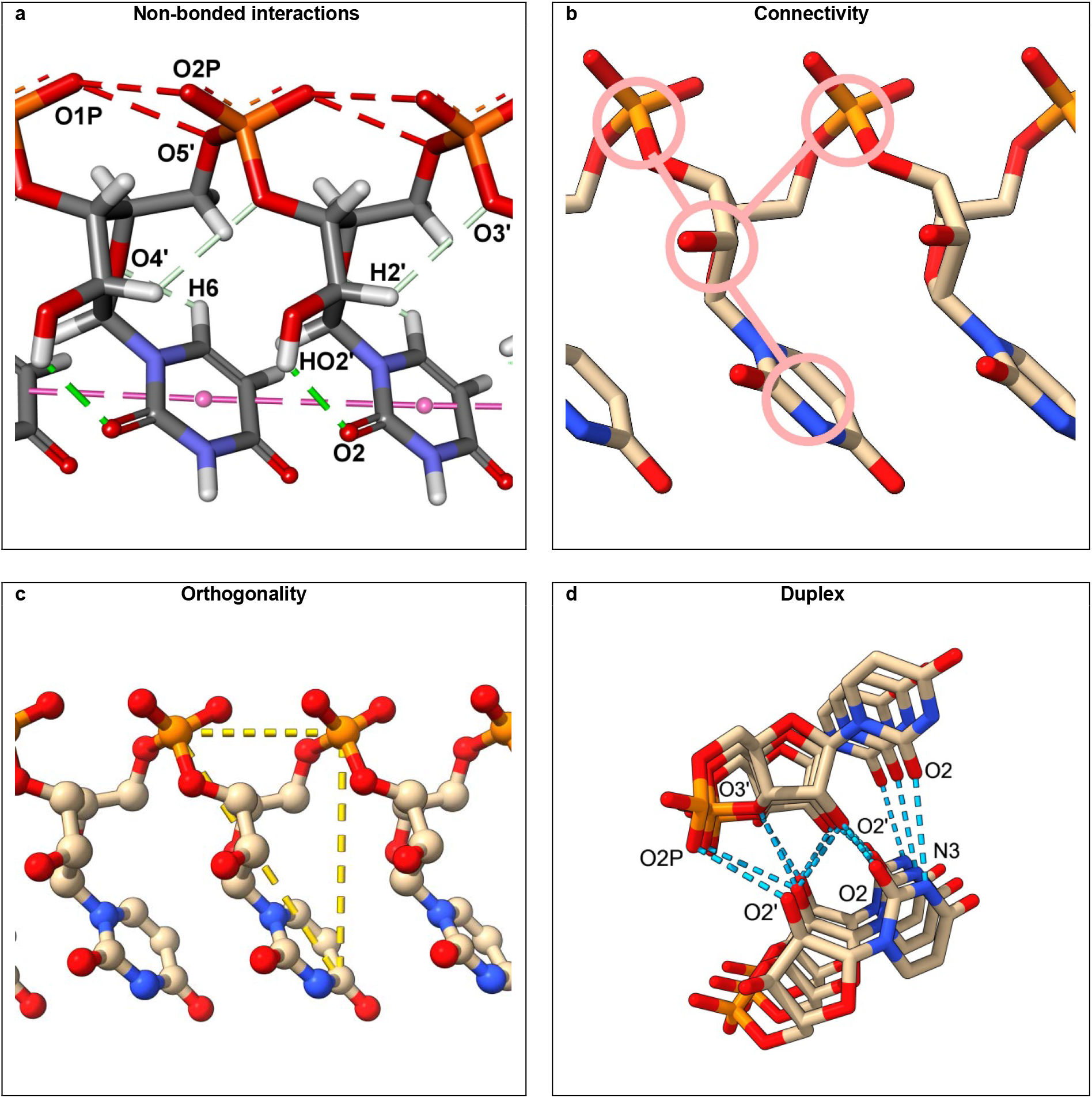
Ortho-RNA. **a**. oRNA is stabilised by internal hydrogen bonds and base stacking. **b.** Connectivity (three-blob pattern) is the same as EDs, *cf*. Extended Data Fig. 2. **c.** oRNA is a remarkably straight molecular. **d.** Two strands of oRNA may form a duplex, linked by hydrogen bonds. Image **(a)** created with BIOVIA, Dassault Systèmes, Discovery Studio Visualizer, v21.1.0.20298, San Diego, Dassault Systèmes, 2021, other images created with UCSF ChimeraX^45^, connectivity overlay created with ImageJ^54^.

**Extended Data Fig. 5.**
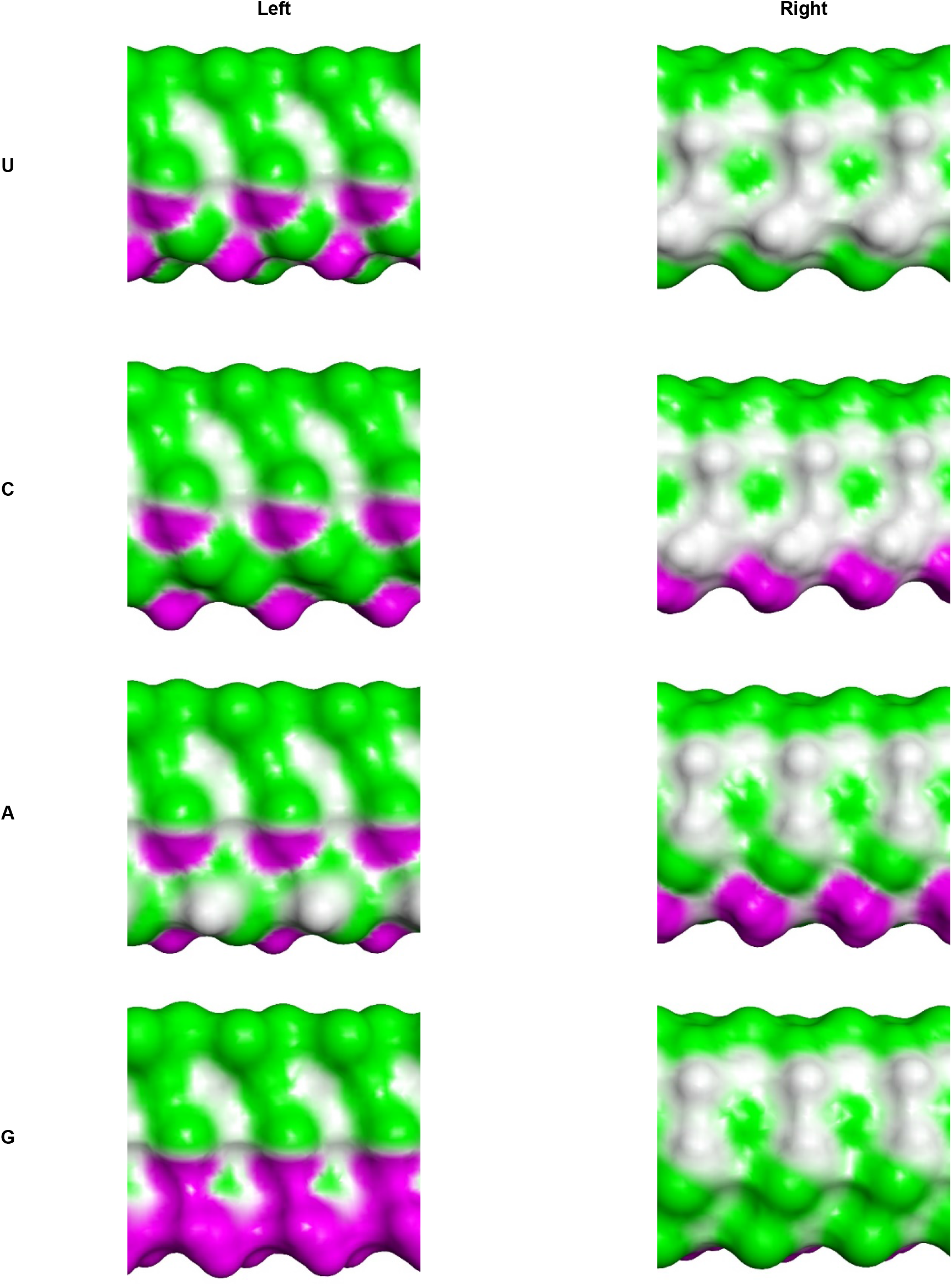
Molecular barcodes. Surfaces of oRNAs (polyU, polyC, polyA and polyG), showing projection of hydrogen-bonding acceptor atoms (green), donor atoms (purple) and non-hydrogen-bonding atoms (white), resembling barcodes readable by the protein environment. Images created with BIOVIA, Dassault Systèmes, Discovery Studio Visualizer, v21.1.0.20298, San Diego, Dassault Systèmes, 2021.

**Extended Data Fig. 6.**
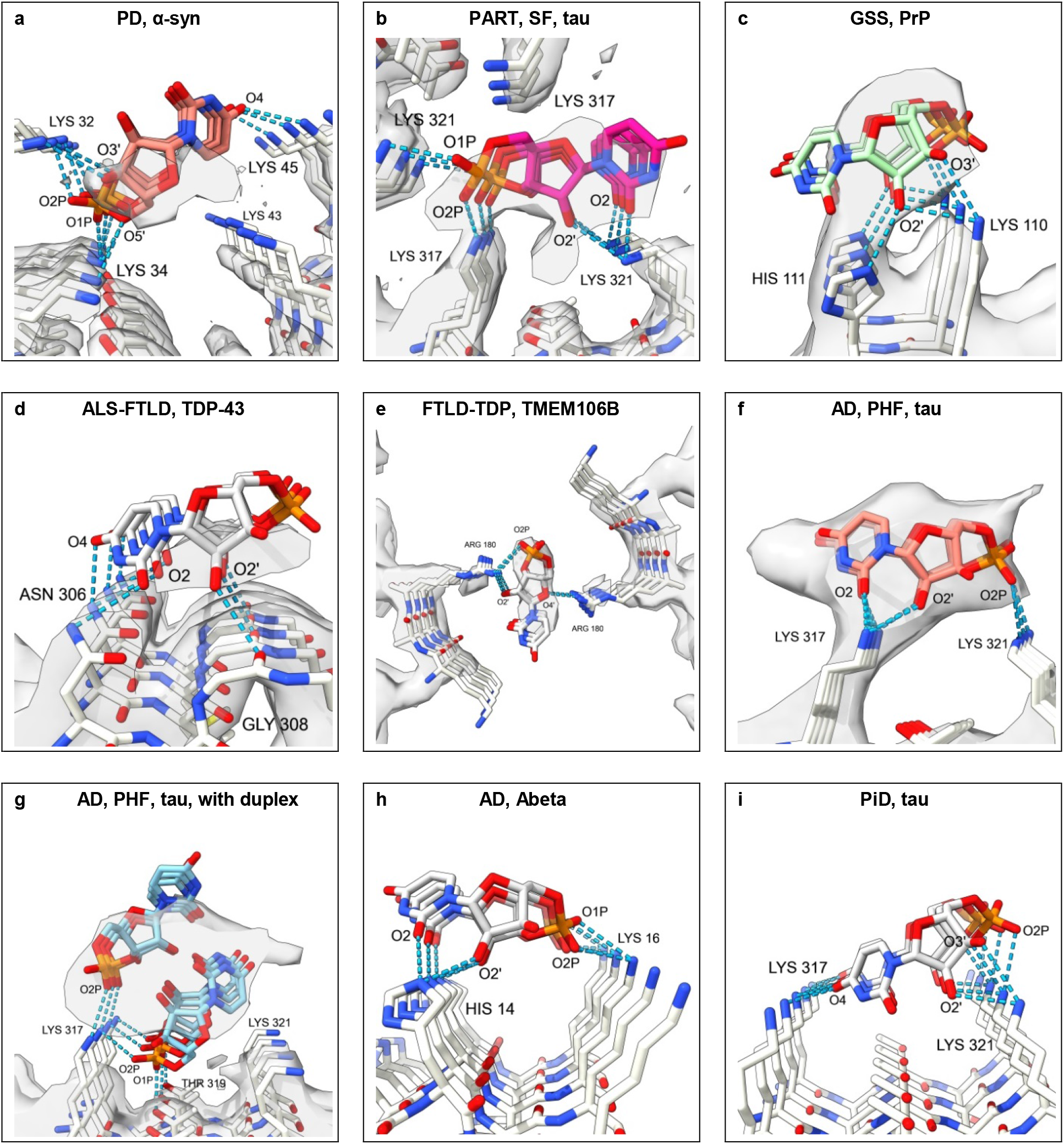
Organising factor. Ortho-RNA forms a rich symmetrical network of hydrogen bonds with proteins, consistent with an organising role in neurodegeneration. Images of EMDBs and PDBs 15285 and 8a9l^6^ **(a)**, 12550 and 7nrs^4^ **(b)**, 26613 and 7un5^18^ **(c)**, 13708 and 7py2^7^**(d)**, 24954 and 7sar^14^ **(e)**, 3741 and 5o31^1^**(f)**, 3742 and 5o3o^1^ **(g)**, 7q4b^37^ **(h)** and 6gx5^8^ **(i)**created with UCSF ChimeraX^45^, oRNA poses generated with AutoDock Vina^36^.

**Extended Data Fig. 7.**
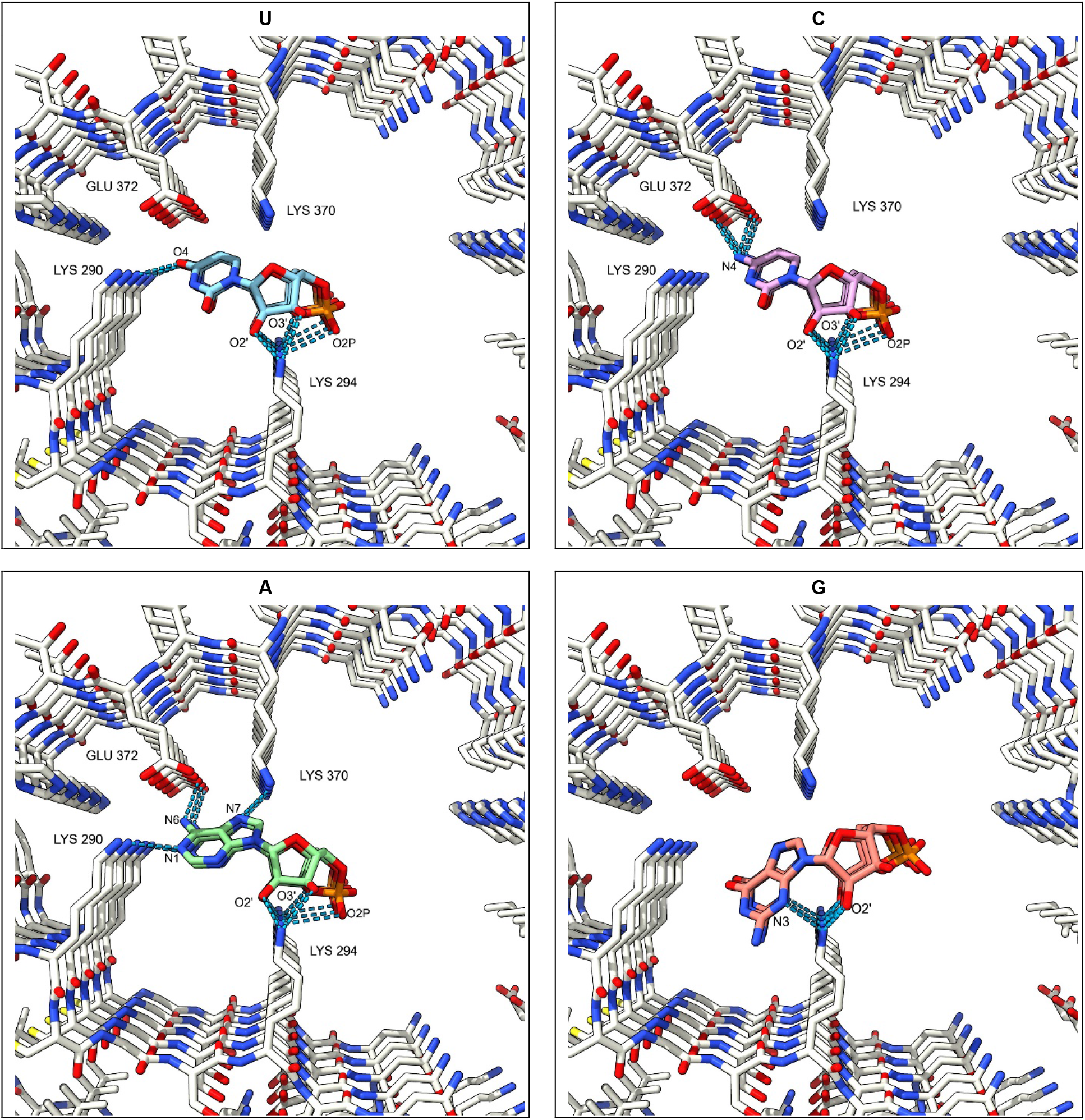
RNA sequence as determinant of fibril structure. oRNAs (polyU, polyC, polyA and polyG) form distinct patterns of hydrogen bonds with K290, K294, K370 and E372 in tau fibril of AGD, consistent with RNA sequence as determinant of fibril structure. Protein model sourced from PDB 7p6d^13^, poses generated with AutoDock Vina^36^, images created with UCSF ChimeraX^45^.

**Extended Data Fig. 8.**
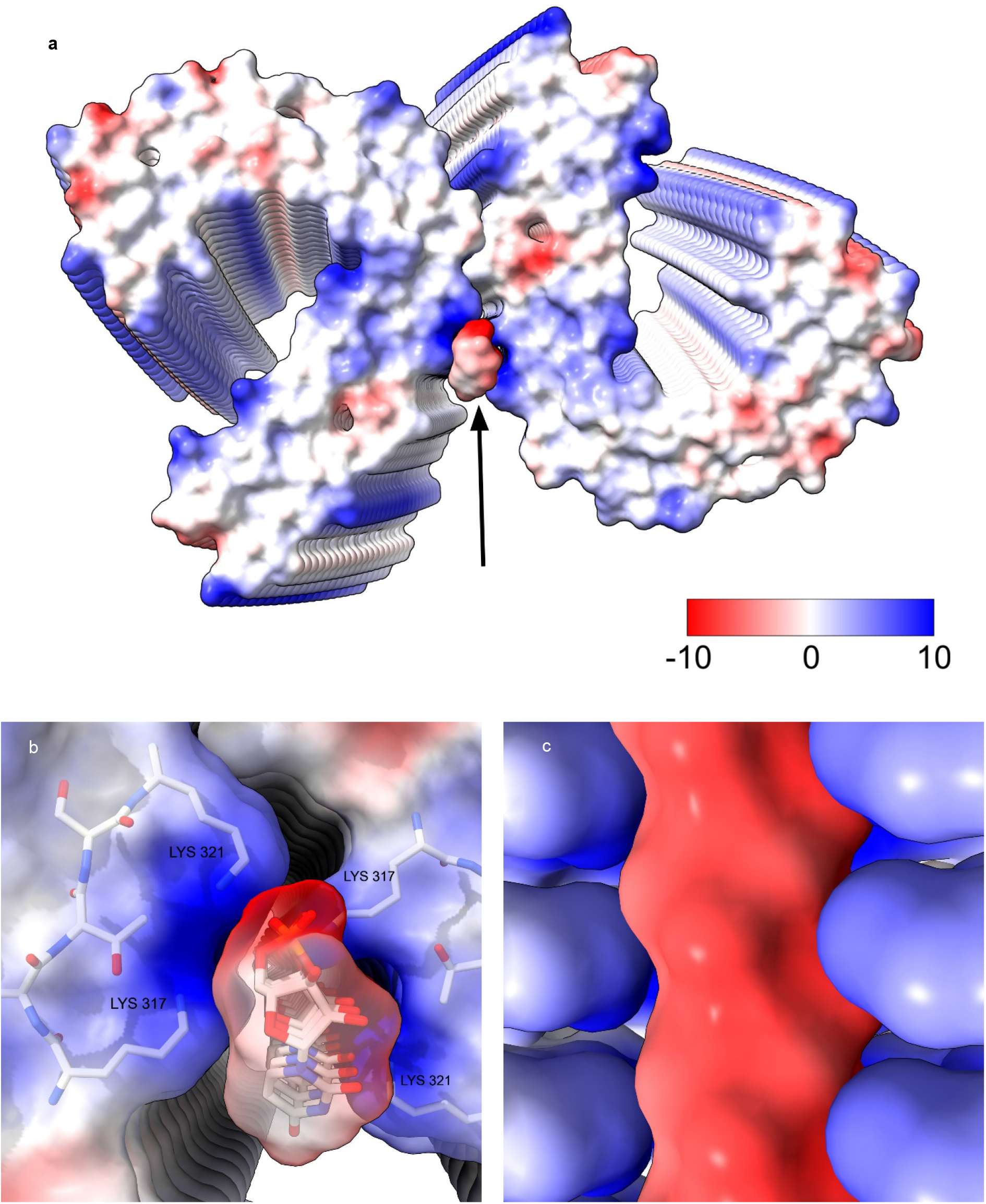
Attraction of opposites. Electrostatic potential surfaces of ortho-RNA docked to protein. **a.** oRNA (arrow) between protofilaments of tau straight filament of PART (units, kcal/mol). **b.** Electrostatic potential surfaces (transparent) and underlying atoms of oRNA and K317-K321. **c.** oRNA (electronegative, red, parallel to fibril axis) between lysines (electropositive, blue, perpendicular to fibril axis), undersurface of fibril, viewed in direction of arrow. Image of PDB 7nrs^4^ created with UCSF ChimeraX^45^, oRNA pose generated in AutoDock Vina^36^.

**Extended Data Fig. 9.**
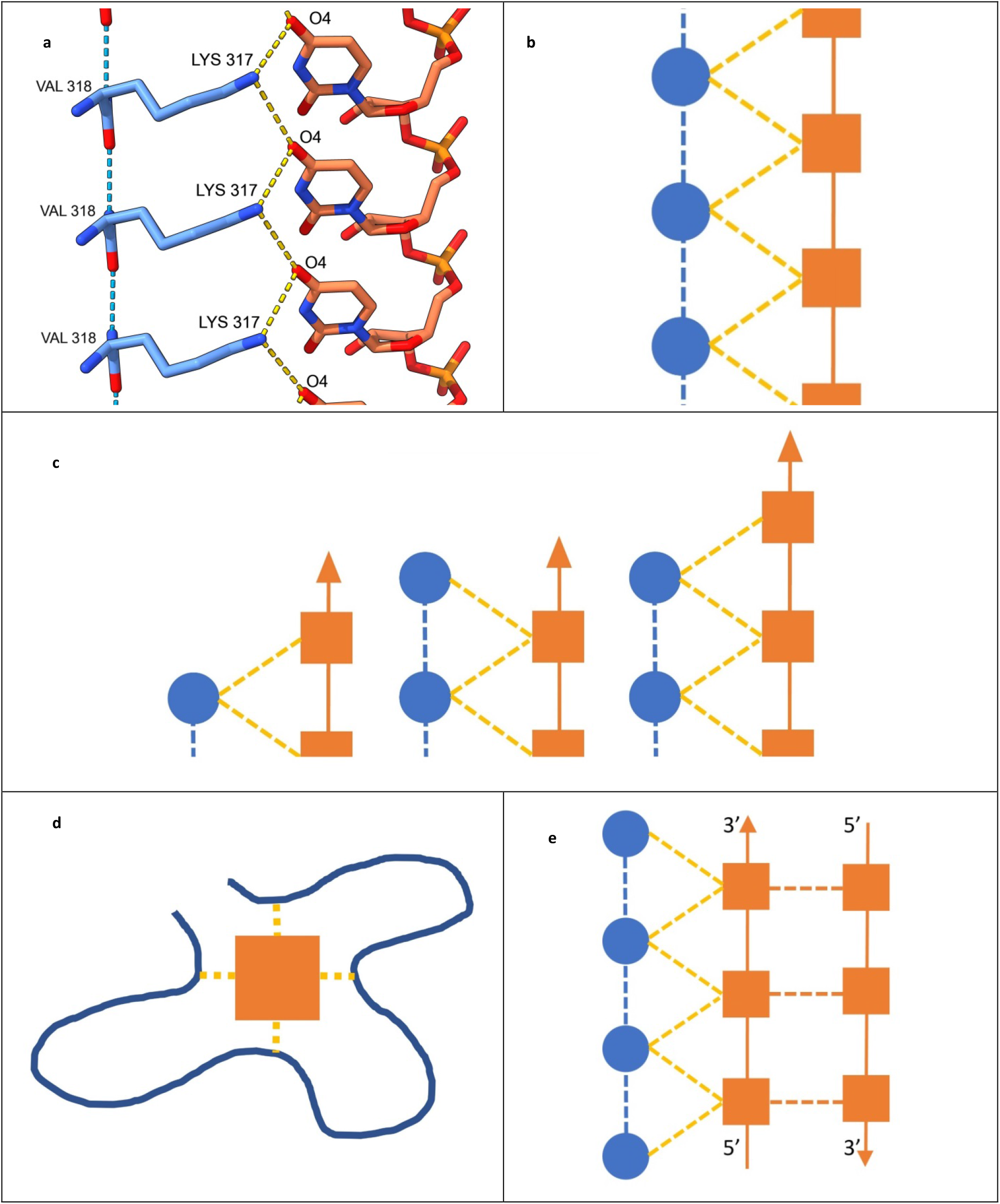
Protein ortho-nucleic acid complex (ponc). **a.** oRNA docked to tau fibril of Pick’s disease (axis runs vertically). **b.**Fibril diagram (corresponds to **a.**). **c.** Proposed mechanism of fibril growth. **d.** Proposed mechanism of protein folding. **e.** Proposed mechanism of fibril replication. Protein rungs (circles), nucleotides (squares), hydrogen bonds (dotted lines), covalent bonds (solid lines); in **d.** protein is solid line. Image of PDB 6gx5^8^ created with UCSF ChimeraX^45^, oRNA pose generated in AutoDock Vina^36^ **(a)**, diagrams **(b-d)** drawn in Microsoft Word.

**Extended Data Table 1.** Estimated molecular weights and docking scores.

| **ED** | **10650-43** | **15285-32** | **12550-317** | **26613-110** | **13708-306** | **24954-180** | **3741-317** | **3742-317** |
| --- | --- | --- | --- | --- | --- | --- | --- | --- |
| Reference | Schweighauser *et al.* 2020<sup>11</sup> | Yang *et al.* 2022<sup>6</sup> | Shi *et al.* 2021<sup>4</sup> | Hallinan *et al.* 2022<sup>18</sup> | Arseni *et al.* 2022<sup>7</sup> | Jiang *et al.* 2022<sup>14</sup> | Fitzpatrick *et al.* 2017<sup>1</sup> | Fitzpatrick *et al.* 2017<sup>1</sup> |
| Disease | MSA | PD | PART | GSS | ALS-FTLD | FTLD-TDP | AD | AD |
| Protein | α-syn | α-syn | tau | PrP | TDP-43 | TMEM-106B | tau | tau |
| PDB | 6xyo | 8a9l | 7nrs | 7un5 | 7py2 | 7sar | 5o3l | 5o3o |
| EMDB | 10650 | 15285 | 12550 | 26613 | 13708 | 24954 | 3741 | 3742 |
| First coordinating residue | K43 | K32 | K317 | K110 | N306 | R180 | K317 | K317 |
| Occupancy | High | Low | High | High | Low | High | High | High |
| Molecular weight (Da) | 275 | 81 | 276 | 360 | 51 | 328 | 309 | 580 |
| AutoDock Vina<sup>36</sup> score (kcal/mol) | -8.9 | -8.7 | -10 | -7.7 | -7.5 | -7.9 | -6 | -9.7 |
| USCF ChimeraX<sup>45</sup> correlation | 0.7 | 0.6 | 0.74 | 0.76 | 0.58 | 0.77 | 0.81 | 0.74 |
| MolProbity<sup>35</sup> clashscore | 2.19 | 8.16 | 4.3 | 0 | 0 | 9.36 | 0 | 1.66 |

